# Neural Basis of Acoustic Species Recognition in a Cryptic Species Complex

**DOI:** 10.1101/2021.08.25.457722

**Authors:** Saumya Gupta, Rishi K. Alluri, Gary J. Rose, Mark A. Bee

## Abstract

Sexual traits that promote species recognition are important drivers of reproductive isolation, especially among closely related species. Identifying neural processes that shape species differences in recognition is crucial for understanding the causal mechanisms of reproductive isolation. Temporal patterns are salient features of sexual signals that are widely used in species recognition by several taxa, including anurans. Recent advances in our understanding of temporal processing by the anuran auditory system provide an excellent opportunity to investigate the neural basis of species-specific recognition. The anuran inferior colliculus (IC) consists of neurons that are selective for temporal features of calls. Of potential relevance are auditory neurons known as interval-counting neurons (ICNs) that are often selective for the pulse rate of conspecific advertisement calls. Here, we took advantage of a species differences in temporal selectivity for pulsatile advertisement calls exhibited by two cryptic species of gray treefrog (*Hyla chrysoscelis* and *Hyla versicolor*) to test the hypothesis that ICNs mediate acoustic species recognition. We tested this hypothesis by examining the extent to which the threshold number of pulses required to elicit behavioral responses from females and neural responses from ICNs was similar within each species but potentially different between the two species. In support of our hypothesis, we found that a species difference in behavioral pulse number thresholds corresponded closely to a parallel species difference in neural pulse number thresholds. However, this relationship held only for ICNs that exhibited band-pass tuning for conspecific pulse rates. Together, these findings suggest that differences in temporal processing of a subset of ICNs provide a mechanistic explanation for reproductive isolation between two cryptic and syntopically breeding treefrog species.

**Summary Statement:** Temporal processing by a subset of midbrain auditory neurons plays key roles in decoding information about species identity in anurans.

## INTRODUCTION

Members of closely related species often share the same ecological niches and exhibit similar phenotypes (Burns and Strauss, 2011; Gholamhosseini et al., 2013; Harper et al., 1961). Consequently, they are more likely to interbreed and produce hybrids, which typically have lower fitness than the two parental species (Gröning and Hochkirch, 2008; Pfennig and Simovich, 2002). Mechanisms that promote premating reproductive isolation are, therefore, crucial for minimizing hybridization-related fitness costs (Coyne and Orr, 2004; Pfennig and Pfennig, 2009). In many animals, especially closely related species living in sympatry, species-specific sexual signals act as a strong premating isolation mechanism. It is often the case that differences in sexual signals of otherwise morphologically and behaviorally similar species are sufficient to facilitate species recognition (Bailey et al., 2017; Couldridge and Alexander, 2002; González-Rojas et al., 2020; Kitano et al., 2008; Miles et al., 2018; Winters et al., 2020).

Animals frequently convey information about their species identity using complex sexual advertisement signals in which different signal elements are produced in distinct temporal patterns (e.g., pulses, notes, syllables, motifs) (Gerhardt and Huber, 2002; Pollack and Hoy, 1979). The temporal patterns of sexual signals often differ among closely related species across a diversity of taxa, including insects (Eberhard and Picker, 2008; Forrest et al., 2006; Jang and Gerhardt, 2006), anurans (Lemmon, 2009; Littlejohn, 1965; Tobias et al., 2011), electric fish (Arnegard et al., 2006; Feulner et al., 2009), and birds (Seneviratne et al., 2012; Wilkins et al., 2018). Many animals, thus, exploit differences in temporal patterns to behaviorally discriminate between signals produced by potential mates of their own species and signalers of other species (Bailey et al., 2017; Deily and Schul, 2004; Gerhardt, 1994; Kollarits et al., 2017; Symes, 2018). Such behavioral selectivity is often mirrored by the selectivity of temporally sensitive neurons (Carlson, 2009; Clemens et al., 2018; Gerhardt and Huber, 2002; Ronacher and Stumpner, 1988). However, differences in the elements of neural circuits that are associated with species differences in recognition remain much less known (but see Triblehorn and Schul, 2009 and Vélez et al., 2017).

Many anurans use temporal sound patterns for species recognition, thus providing an excellent opportunity to investigate the neural basis of species differences in temporal pattern recognition. The advertisement calls male anurans produce to attract females frequently consist of repeated pulses with species-specific shapes and durations that are produced at species-specific pulse rates (Brown and Brown, 1972; Gerhardt and Doherty, 1988; Littlejohn, 1965; Penna and Veloso, 1990; Vélez et al., 2012). Behavioral studies have shown that female anurans are highly selective for temporal features that correspond to conspecific advertisement calls (Bush et al., 2002; Gerhardt, 1989; Kruse, 1981; Loftus-Hills and Littlejohn, 1971; Penna, 1997). Parallel neurophysiological studies of auditory processing have further shown that the central auditory system of anurans consists of an ensemble of neuronal filters that are selective for distinct temporal features of calls (Feng et al., 1990; Hall, 1994; Rose, 1995; Rose and Capranica, 1983). One potentially important class of auditory neuron in the anuran inferior colliculus (IC; homolog of the torus semicircularis) are the ‘interval-counting neurons’ (ICNs), which show species-specific tuning in pulse rate, an important species recognition cue in anurans (Gerhardt and Huber, 2002; Schul and Bush, 2002; Schwartz, 1987). These cells respond with action potentials only after a threshold number of sound pulses are presented with inter-pulse-onset intervals that correspond to species-typical pulse rates (Alder and Rose, 1998; Edwards et al., 2002). Even a single interval that is shorter or longer than the optimal range can reset the interval-counting process (Edwards and Rose, 2003; Edwards et al., 2002). Since the discovery of these neurons, several behavioral studies have suggested that ICNs might be involved in call recognition (Rose and Brenowitz, 2002; Schwartz et al., 2010; Vélez and Bee, 2011). But whether species-differences in mate choice decisions involving conspecific recognition in anurans reflect species-differences in the activity of ICNs remains to be investigated.

In this study, we took advantage of a natural experiment of divergence in temporal sound pattern recognition associated with polyploid speciation in gray treefrogs to test the hypothesis that ICNs mediate acoustic species recognition. The gray treefrogs form a cryptic species complex consisting of two closely related species, the diploid Cope’s gray treefrog, *Hyla chrysoscelis*, and the tetraploid eastern gray treefrog, *Hyla versicolor*. Current evidence suggests that *H. versicolor* evolved as a result of autopolyploidization from a now extinct lineage of *H. chrysoscelis* (Bogart et al., 2020; Booker et al., 2020). The two species are morphologically indistinguishable and breed at the same times and places across much of their shared geographic range. Males of both species produce pulsatile advertisement calls that are highly similar spectrally but differ in their temporal patterns (Fig. 1) (Gerhardt, 1994; Gerhardt, 2005). Compared with the calls of *H. chrysoscelis*, the calls of *H. versicolor* have slower pulse rates and consist of longer pulses that rise more slowly in amplitude (Gerhardt and Doherty, 1988). Females of both species rely on species-specific temporal patterns to discriminate between the calls of conspecific and heterospecific males. Female *H. chrysoscelis* prefer calls with a conspecific pulse rate of 30-60 pulses/s over calls with slower and faster pulse rates, whereas females of *H. versicolor* prefer calls with longer pulses and slower pulse rise times and that are produced at a slower pulse rate of 10-30 pulses/s (Bush et al., 2002; Gerhardt and Doherty, 1988). The slower pulse-rate preference of female *H. versicolor* appears to be a byproduct of their selectivity for pulses that have slower rise and longer duration (Schul and Bush, 2002).

**Fig. 1.**
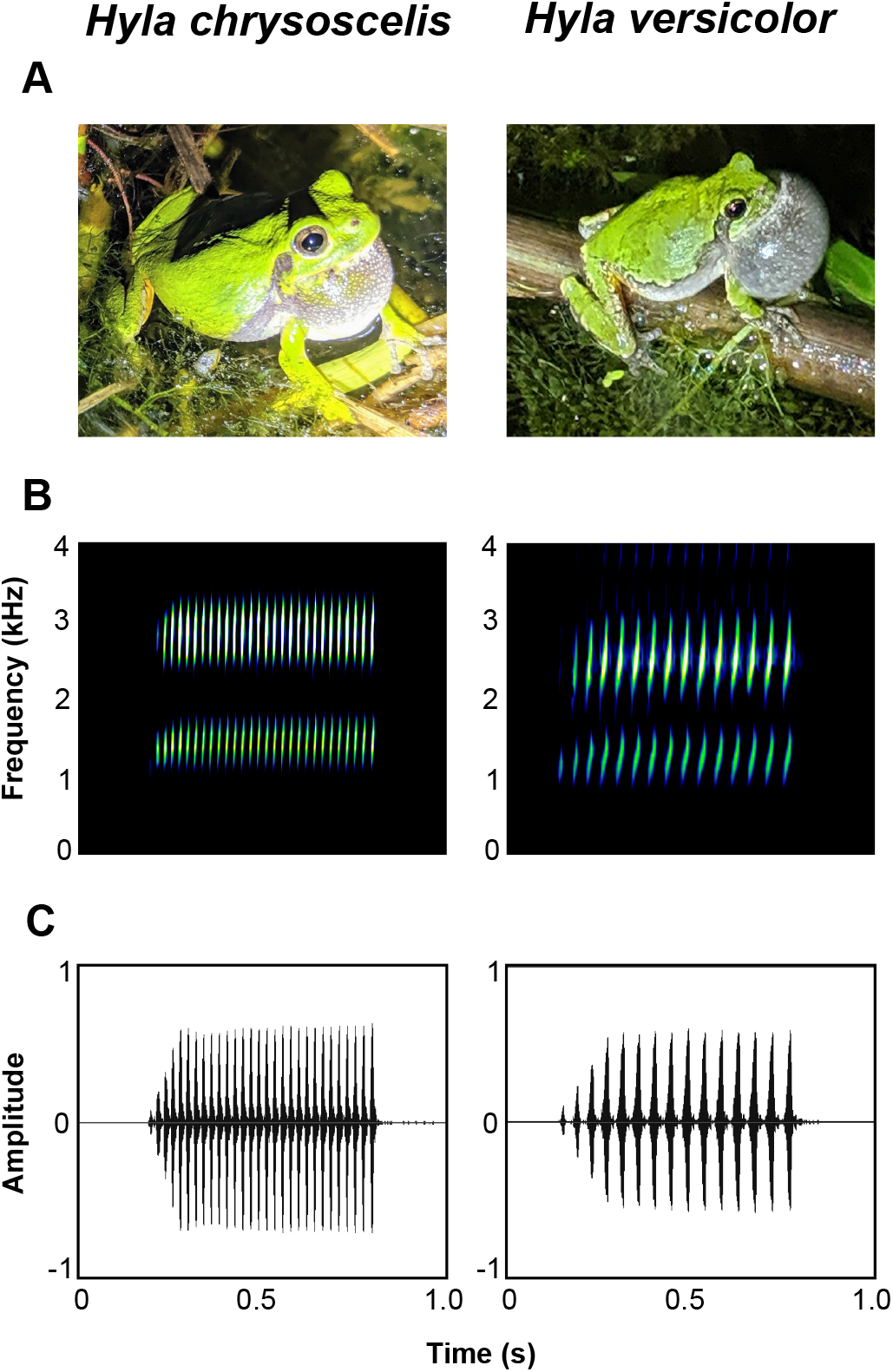
Advertisement calls of the two visually indistinguishable species of gray treefrogs. (A) Pictures of calling males of *Hyla chrysoscelis* and *Hyla versicolor*. (B) Spectrograms (plot of frequency vs. time) depicting the pulsatile structure and similar bimodal frequency spectrum of the species-specific advertisement calls. (C) Oscillograms (plot of amplitude vs. time) depicting species difference in pulse rates.

Neurons in the gray treefrog IC exhibit patterns of pulse-rate and pulse rise-time selectivity similar to those exhibited by females choosing mates, suggesting that temporal selectivity of midbrain neurons contributes to the selectivity of behavioral responses that mediate mating decisions (Hanson et al., 2016; Rose et al., 1985; Rose et al., 2015). In addition, both ICNs and female gray treefrogs only respond to conspecific advertisement calls that exceed a threshold number of pulses in duration (Bush et al., 2002; Edwards et al., 2002; Rose et al., 2015; Vélez and Bee, 2011). However, the degree of correspondence between behavioral and neural pulse number thresholds remains unknown. Investigating this correspondence *within* and *between* the two gray treefrog species could shed light on the neural basis of species-specific recognition. According to the hypothesis that ICNs mediate acoustic species recognition, our primary prediction was that the threshold number of pulses required to elicit behavioral responses from females and neural responses from ICNs would be similar within each species but potentially different between species. To test this prediction, we used an adaptive tracking procedure (Bee and Schwartz, 2009) to determine the minimum number of pulses produced at conspecific pulse rates required to elicit positive phonotaxis from females of each species. We then used single-unit, extracellular recordings and whole-cell recordings to determine neural pulse number thresholds of ICNs in the two species, focusing specifically on two populations of ICNs that exhibit band-pass versus band-suppression tuning for pulse rate (Edwards and Rose, 2003; Rose, 2014).

## MATERIALS AND METHODS

### Behavioral Pulse Number Thresholds

#### Animals

Our protocols for collecting and handling females are described in more detail elsewhere (Gupta and Bee, 2020; LaBarbera et al., 2020). Briefly, we collected 44 gravid female *H. chrysoscelis* and *H. versicolor* in amplexus from a sympatric population at the Tamarack Nature Center (Ramsey County, MN, USA) during the 2019 breeding season (May – June). All collections were made at night between 2200-0100 h. Each collected pair was put in a separate small plastic container and transported back to the laboratory on the St. Paul campus of the University of Minnesota, where behavioral tests were conducted. We provided pairs with aged tap water and maintained them at 4°C prior to testing to postpone egg-laying and maintain female responsiveness (Gerhardt, 1995). We tested subjects within 72 hours of collection and promptly released them back at the collection site at the completion of testing.

#### Experimental Setup and Acoustic Stimuli

Positive phonotaxis by female frogs indicates the recognition of an acoustic stimulus as the advertisement call of a potential conspecific mate (Gerhardt and Huber, 2002; Ryan and Rand, 2001). Hence, we used phonotaxis as an assay to determine behavioral pulse number thresholds. The experimental setup for conducting phonotaxis tests has been described in previous studies of signal recognition and discrimination (Bee and Schwartz, 2009; LaBarbera et al., 2020; Nityananda and Bee, 2012; Vélez and Bee, 2013), and readers are referred to those studies for additional details not reported here. Briefly, a circular phonotaxis arena (2-m diameter, 60-cm height) was set up inside a temperature-controlled (20 ± 1 °C) hemi-anechoic sound chamber (length × width × height: 2.8 × 2.3 × 2.1 m; Industrial Acoustics Company, IAC, North Aurora, IL, USA). The inside walls and ceiling of the sound chamber were lined with dark gray acoustic absorber panels (IAC’s Planarchoic^™^ system), and the flooring consisted of dark gray, low-pile carpet. The arena itself was constructed from hardware cloth covered with black fabric. An acoustically transparent cage (9-cm diameter, 2-cm height), from which subjects were released at the beginning of each test, was located at the center of the arena floor. The lid of the release cage could be lifted remotely via a pulley system. Just outside the arena wall, two speakers separated by 90° were located on the floor and directed towards the release cage. A response zone consisting of a 10-cm semi-circle was marked on the arena floor in front of each speaker. Movements of subjects inside the arena were observed under infrared (IR) illumination and responses were scored in real-time using a video monitor located outside the chamber.

Acoustic stimuli (44.1 kHz, 16 bit) were synthesized in MATLAB 2018b (Mathworks, Natick, MA, USA). All stimuli consisted of a sequence of advertisement calls designed to simulate a calling male. Calls consisted of a sequence of pulses and were modeled after the natural advertisement calls produced by males of *H. chrysoscelis* (Ward et al., 2013) and *H. versicolor* males (Tables S1,S2), recorded at our study sites, with spectral and temporal properties adjusted to a temperature of 20°C following Platz and Forester (1988). For *H. chrysoscelis*, each pulse was 10 ms in duration (3.1-ms inverse exponential rise time; 5.4-ms exponential fall time) and was created by adding two phase-locked and harmonically related sinusoids with frequencies (and relative amplitudes) of 1.25 kHz (-11 dB) and 2.5 kHz (0 dB). The inter-pulse interval, defined as the duration between the onsets of two consecutive pulses, was 20 ms (50 pulses/s, 50% duty cycle). The pulses composing *Hyla versicolor* calls had a duration of 30 ms (20-ms linear rise time; 10-ms linear fall time) and were created by adding two phase-locked and harmonically related sinusoids with frequencies (and relative amplitudes) of 1.2 kHz (-5 dB) and 2.4 kHz (0 dB). The inter-pulse interval was 60 ms (16.67 pulses/s, 50% pulse duty cycle).

Because females of the two cryptic species were collected from a sympatric population, the first phonotaxis test was always a species identification test that consisted of repeatedly alternating a *H. chrysoscelis* call with a *H. versicolor* call from the two speakers, with each call being average length, repeated at a rate of 11 calls/min, and preceded and followed by equivalent intervals. Each gray treefrog species is highly selective for the pulse rates of conspecific calls and discriminates against pulse rates of the other species’ calls (Bush et al., 2002; Gerhardt and Doherty, 1988; Gupta and Bee, 2020). Therefore, we determined each subject’s species identity based on the call alternative chosen in this initial species identity test. We subsequently performed a set of no-choice tests to measure pulse number thresholds. The stimulus for each no-choice test consisted of a sequence of advertisement calls having temporal properties specific to the appropriate species with the single exception of the number of pulses per call. We manipulated the number of pulses per call across stimuli in different no-choice tests but held it constant within a particular stimulus sequence used in each no-choice test. Within a stimulus, calls were presented at species-typical call rates (11 calls/min for *H. chrysoscelis* and 10 calls/min for *H. versicolor*).

All stimuli were broadcast using a playback system consisting of Adobe Audition 3.0 (Adobe Systems Inc., San Jose, CA, USA) that interfaced with a MOTU model 16A sound card (MOTU, Inc., Cambridge, MA, USA) and Crown XLS 1000 High-Density Power Amplifiers (Crown International, Los Angeles, CA, USA). The final output signals were broadcast through Mod1 Orb speakers (Orb Audio, Sherman Oaks, CA, USA). The frequency response of the playback system was flat (± 2.5 dB) across the frequency range of interest. Sound pressure levels (SPL re 20 μPa, fast, C-weighted) were calibrated by placing a Bruёl and Kjær Type 4950 microphone connected to a Bruёl and Kjær Type 2250-L sound level meter (Bruёl and Kjær, Nærum, Denmark) at the approximate position of a subject’s head while sitting in the release cage. In different trials, stimuli were calibrated to SPLs of 85 dB or 65 dB (at 1 m) to investigate the level-dependence of behavioral pulse number thresholds, as auditory mechanisms in frogs can be level-dependent (Gerhardt, 2005; Gerhardt, 2008). We used the 85-dB playback level for the species identification test because a sound pressure level of 85 dB approximates the amplitude of a male calling at a distance of 1 m (Gerhardt, 1975).

#### Experimental Protocol

Every phonotaxis test was conducted using the same protocol. Approximately 40 min prior to commencing tests with a given subject, the female and her chosen mate were transferred to an incubator and allowed to reach a body temperature of 20 ± 1°C. Before the start of each test, the subject was separated from her mate and placed in the release cage. A test began with a 30-s period of silence followed by two call repetitions of the test stimulus, after which subjects were remotely released from the cage and allowed to respond. After the initial species identification test, each subject’s pulse number threshold was determined at one of the two signal amplitudes and then at the second one, with the order randomly determined for each subject.

We operationally defined behavioral pulse number thresholds as the minimum number of pulses required to elicit positive phonotaxis in a no-choice test. We defined positive phonotaxis as occurring when the subject exhibited directed hopping or walking movements and entered the response zone in front of the active speaker. An adaptive tracking procedure developed to measure signal recognition thresholds (Bee and Schwartz, 2009) was followed to measure behavioral pulse number thresholds. This procedure involved a sequence of no-choice tests in which we systematically varied the number of pulses per call in each test based on the subject’s response in the immediately preceding test. In the initial no-choice test at a given signal amplitude, the stimulus call consisted of either 6 or 8 pulses. In subsequent tests, the pulse number was increased or decreased by 2 pulses based on the subject’s response in the preceding test. For example, if a subject responded to an 8-pulse call, it was presented with 6-pulse call in the next test, but if it failed to respond to an 8-pulse call, it was presented with a 10-pulse call in the next test. This decrease or increase in pulse number continued until a subject changed its behavior between two consecutive tests, going either from a no response to a response or from a response to a no response. In the next and final test, the number of pulses in the stimulus call was either decreased or increased by 1 pulse depending, respectively, on whether or not the subject responded in the previous test. If the subject did not respond in the final test, a ‘reference test’ was performed in which a sequence of average-length conspecific calls was presented to verify if the subject was still motivated to respond. One subject did not respond in this reference test during its threshold measurement at 85 dB SPL, and hence, that threshold was removed from statistical analyses. We removed another subject’s threshold at 65 dB SPL because of a procedural error made during its threshold determination. The pulse number threshold of each subject at each signal amplitude was computed by taking the mean of the lowest pulse number that elicited a response and the highest pulse number that failed to do so.

### Neural Pulse Number Thresholds

#### Animals

Our protocols for conducting electrophysiological experiments to determine pulse number thresholds are described in more detail elsewhere (Hanson et al. 2015; Rose et al. 2015), and readers are referred to those studies for details not presented here. We used 14 wild-caught female *H. chrysoscelis* and *H. versicolor* from central Minnesota as subjects for single-unit extracellular and whole-cell recordings. Subjects were group-housed at the University of Utah in 30 × 60 × 30 cm glass tanks that were kept inside a room with a 12-hour light: dark cycle. Subjects were provided with *ad libitum* access to water and fed crickets twice a week.

#### Surgery

Subjects were anesthetized by immersion in 0.1% MS-222. Following the loss of reflexes, they were loosely wrapped in moist surgical gauze to facilitate cutaneous respiration, and 2% lidocaine hydrochloride was topically applied to the skin on the dorsal surface of the head. A craniotomy was performed to make a small hole in the skull to expose the optic tectum. The hole was subsequently filled with Gelfoam (Ethicon Inc., Bridgewater, NJ, USA), and subjects were allowed to recover overnight.

#### Stimulus Generation and Presentation

We synthetically constructed all acoustic stimuli (24144.0625 Hz sample rate) using an auditory stimulus generator (Tucker Davis Technologies, Alachua, FL, USA) and custom software written in MATLAB 2011b. Since most IC neurons do not respond to pure tones, we used amplitude modulated stimuli (Alder and Rose, 2000). Sinusoidal amplitude-modulated (SAM) stimuli were generated by multiplying a pure tone with a sinusoidal modulating waveform. Stimuli consisting of pulses of natural shape were generated by multiplying a pure tone with a modulating envelope that was a mathematical representation of the natural pulse envelope of either *H. chrysoscelis* or *H. versicolor* calls. More details on how these sounds were generated are described in Alder and Rose (2000).

We amplified (SA1; Tucker-Davis Technologies, Alachua, FL, USA) and presented each stimulus free-field in the sound chamber from one of two Bose speakers (Model 101; Bose corporation, Framingham, MA, USA) that were placed 0.5 m to the left or right of the subject and pointed directly towards the subject’s tympanum. The active speaker was always the one that was contralateral to the recording site. We measured sound pressure levels using an IE-30A sound level meter (Ivie Technologies, Inc., Springville, UT, USA) connected to a microphone (ACO Pacific, Inc., Belmont, CA, USA) and Cetec Ivie IE-2P preamp (Ivie Technologies, Inc., Springville, UT, USA) that was situated just above the subject.

#### Electrode Construction

We made microelectrodes from borosilicate glass capillary tubes (1mm outer diameter, 0.58 mm inner diameter; A-M systems #5960, Sequim, WA, USA) using a Flaming-Brown type puller (model P-97; Sutter Instruments, Novato, CA, USA). The extracellular electrodes had tip diameters of 2-3 μm and resistances between 0.8 and 1.2 MΩ when filled with 2 M NaCl solution. Patch pipettes for whole-cell recordings had an outside tip diameter of approximately 1-1.5 μm and resistances between 10 and 20 MΩ. The tips of these pipettes were backfilled with a solution (pH = 7.4) consisting of 100 mM potassium gluconate, 2 mM KCl, 1 mM MgCl_2_, 5 mM EGTA, 10 mM HEPES, 20 mM KOH, and 20 mM biocytin. We used another solution with similar composition but with 20 mM mannitol instead of the 20 mM biocytin to fill the pipette shanks.

#### Recording Procedure

On the day of electrophysiological recordings, we immobilized the subject with an intramuscular injection of pancuronium bromide (4 μg/g) and wrapped it in moist gauze. The protocols for making neurophysiological recordings were similar to those described elsewhere (Alder and Rose, 2000; Edwards et al., 2007; Rose and Fortune, 1996; Rose et al., 2015). Briefly, the subject was placed on a platform mounted on a vibration isolation table (Technical Manufacturing Company, Peabody, MA, USA) inside a temperature-controlled (20 ± 2°C) hemi-anechoic sound chamber (length × width × height: 2.1 × 2.2 × 2 m; Industrial Acoustics Company, North Aurora, IL, USA). The electrodes were positioned stereotaxically and were advanced into the IC remotely by means of a 3-axis Microdrive (model IVM-3000; Scientifica, East Sussex, UK). Auditory neurons were identified using a search stimulus that consisted of a sequence of SAM tones that varied in modulation frequencies (between 10 to 100 Hz) as well as carrier frequencies (between 150 to 2500 Hz). Upon identifying a single unit that responded to the search stimulus, slight suction was applied to improve recordings.

For whole-cell recordings, the patch pipette was advanced in increments of 1.5 μm after reaching the location of the recording. Positive pressure was maintained at the time of the increments, and -0.1 nA square-wave pulses of 500ms were used to monitor resistance. A small increase (10%) in the voltage indicated that contact with the isolated unit was established. Upon contact, negative pressure was applied to increase the seal resistance to gigaohm levels. Finally, a negative current (~ 0.5 nA) was applied to rupture a tiny portion of the cell. Additional details on whole-cell recording procedures can be found elsewhere (Alluri et al., 2016, 2021)

After isolating a single unit, we determined the unit’s best excitatory frequency (BEF) as the carrier frequency that elicited a response at the lowest threshold, where threshold was defined as the sound pressure level at which a unit spiked in response to at least 50% of the stimulus presentations. All subsequent acoustic stimuli used to record from that unit were generated at the unit’s BEF and were broadcast at amplitudes ~10 dB above the unit’s threshold. Across units, the amplitudes of the stimuli ranged between approximately 65 dB and 90 dB SPL. Each unit’s pulse-rate selectivity was established by presenting pulsatile sounds that had a constant pulse duty cycle but that varied in pulse rate (between 5-80 pulses/s). All pulses in these stimuli had species-typical pulse shapes for the species from which recordings were made (i.e., relatively slower rise time pulses for *H. versicolor* and faster rise time pulses for *H. chrysoscelis*). A unit was considered pulse-rate selective if the number of spikes elicited in response to the most preferred and the least preferred pulse rate differed by at least 50%. A pulse-rate selective neuron was further identified as an interval-counting neuron (ICN) if it elicited spikes after a threshold number of consecutive pulses had occurred at any pulse rate within its range of pulse-rate selectivity.

#### Data Acquisition and Processing

Neural recordings were digitized at 10 kHz using a data acquisition interface (Power 1401, Cambridge Electronic Design, Cambridge, UK) and stored and analyzed using the Spike-2 software (Cambridge Electronic Design, Cambridge, UK). We recorded from 22 ICNs whose pulse-rate selectivity overlapped with the species’ conspecific pulse rates (between 10-30 pulses/s for *H. versicolor* and between 30-60 pulses/s for *H. chrysoscelis*). Results from these units were combined with data obtained from 32 additional units described by Rose et al. (2015) that were investigated using the same procedures and met the same criterion of having preferred pulse-rate selectivity that overlapped the pulse rates of corresponding species’ calls. This yielded a final sample size of 54 units considered in the present study. Following Rose et al. (2015), units were classified as either band-pass ICNs or band-suppression ICNs. Band-pass ICNs were selective for a narrow range of pulse rates, outside of which they showed minimal responsiveness. The best pulse rates of band-pass ICNs were distributed between 15-20 pulses/s in *H. versicolor* and between 40-50 pulses/s in *H. chrysoscelis* (Rose et al., 2015). Band-suppression ICNs, in contrast, showed a 50% or greater reduction in the response within an intermediate range of pulse rates below and above which they exhibited robust responsiveness. Band-suppression ICNs typically exhibited a response minimum at approximately 20 pulses/s in *H. chrysoscelis* and ~30 pulses/s in *H. versicolor* (Rose et al., 2015).

To compare behavioral responses from females with neural responses from ICNs, we focused our analyses on neural thresholds measured in response to stimuli with conspecific pulse rates. We operationally defined a neuron’s pulse number thresholds as the minimum number of sound pulses at which it spiked in response to at least 50% of the presentations (Edwards et al., 2007). Thresholds were determined in one of the two ways. For 43 units, thresholds were determined from sound presentations in which pulse number was systematically varied. For the remaining 11 units, we estimated the pulse-number threshold from responses to pulse trains that were 400 ms in duration; threshold was determined as the number of pulses that preceded initiation of spiking, accounting for an approximately 30-ms latency for conduction of activity to the IC. The average neural pulse number thresholds determined using the two methods differed by less than one pulse. Hence, we combined the data obtained from the two measurements to describe distributions of neural pulse number thresholds.

### Statistical Analysis

The primary prediction of our study was that if ICNs mediate acoustic species recognition, then the threshold number of pulses, presented at conspecific pulse rates, required to elicit behavioral responses from females and neural responses from ICNs should be similar within each species but potentially different between the two species. To this end, we first fitted two nested linear mixed-effects models (LMM) to compare the behavioral pulse number thresholds between the two species (*H. chrysoscelis* and *H. versicolor*) and between the two signal amplitudes (65 dB and 85 dB SPL). The first model included the fixed effects of species and signal amplitude as predictor variables and the random effect of subject ID. The second model included an additional species × signal amplitude interaction term. Next, we fitted two nested linear models to compare neuronal pulse number thresholds between the two species and between band-pass and band-suppression ICNs. The first model included only the main effects of species and ICN populations as the predictor variables, and the second model included an additional species × ICN population interaction term. An ANOVA was used to compare each pair of nested models and the simpler model was adopted if no significant difference was found (using a significance criterion of α = 0.05). In analyses of both behavioral and neural pulse number thresholds, we did not find a significant difference between the two nested models with and without the interaction terms (*p* > 0.05). Hence, the simpler model without the interaction term was adopted in each case (see Tables S3 and S4 for ANOVA results). Finally, a linear model was fitted separately for each species to compare the behavioral thresholds to neural thresholds obtained from each ICN population. In this analysis, pulse number threshold was the dependent variable and whether it was determined behaviorally, from a band-pass ICN, or from a band-suppression ICN was the categorical independent variable. This analysis required us to take into account the fact that behavioral pulse number thresholds were determined within subjects at two fixed signal amplitudes (65 and 85 dB SPL) while neural pulse number thresholds were determined between units across a range of amplitudes (65 to 90 dB SPL) based on each unit’s response threshold. Therefore, as a first approach, a mean behavioral threshold was calculated for each subject for these analyses by averaging its behavioral thresholds 65 dB and 85 dB SPL. A significance criterion of α = 0.025 (after Bonferroni correction) was used for these analyses to control for multiple within-species comparisons of behavioral to neural data. As an alternative approach, we also compared neural thresholds with behavioral thresholds determined separately at 65 dB or 85 dB SPL; these analyses yielded qualitatively similar results (see Tables S5 and S6).

All statistical analyses in this study were done in R v.4.0.2 (R core team, 2020). In all analyses, pulse number thresholds were log_10_ transformed to meet the assumptions of normality and homogeneity of variance. All estimates were subsequently back-transformed to report effect sizes.

## RESULTS

### Behavioral Pulse Number Thresholds

Behavioral pulse number thresholds were higher for *H. chrysoscelis* than for *H. versicolor* at both signal amplitudes (Fig. 2). At 65 dB SPL, the mean (± 95% confidence interval) pulse number threshold of *H. chrysoscelis* was 7.98 ± 1.13 at 65 dB SPL (Median = 7.5, IQR = 5.5-9.5, N = 21) and that of *H. versicolor* was 4.86 ± 1.18 at 65 dB SPL (Median = 3.5, IQR = 3.5-6.5, N = 22). At 85 dB SPL, the mean pulse number threshold of *H. chrysoscelis* was 7.50 ± 0.83 (Median = 7.5, IQR = 5.5-8.5, N = 21) and that of *H*. versicolor was 3.41 ± 0.81 (Median = 2.5, IQR = 2.5-3.5, N = 22). The results of LMM showed that on average, the pulse number threshold of *H. chrysoscelis* was significantly higher (by a factor of 2.14; *β* = 0.33, *p* < 0.001; Table 1) than *H. versicolor* after controlling for the effects of signal amplitude. The threshold determined at 65 dB SPL was also significantly higher *β* = 0.09, *p* = 0.013; Table 1) than that determined at 85 dB SPL after controlling for species differences, but only by a factor of 1.23. Averaged over both signal amplitudes, the pulse number threshold of *H. chrysoscelis* was 7.75 ± 0.81 (Median = 7.75, IQR = 6.4-8.5, N = 20) and that of *H. versicolor* was 4.14 ± 0.90 (Median = 3.5, IQR = 2.5-5.0, N = 22).

**Fig. 2.**
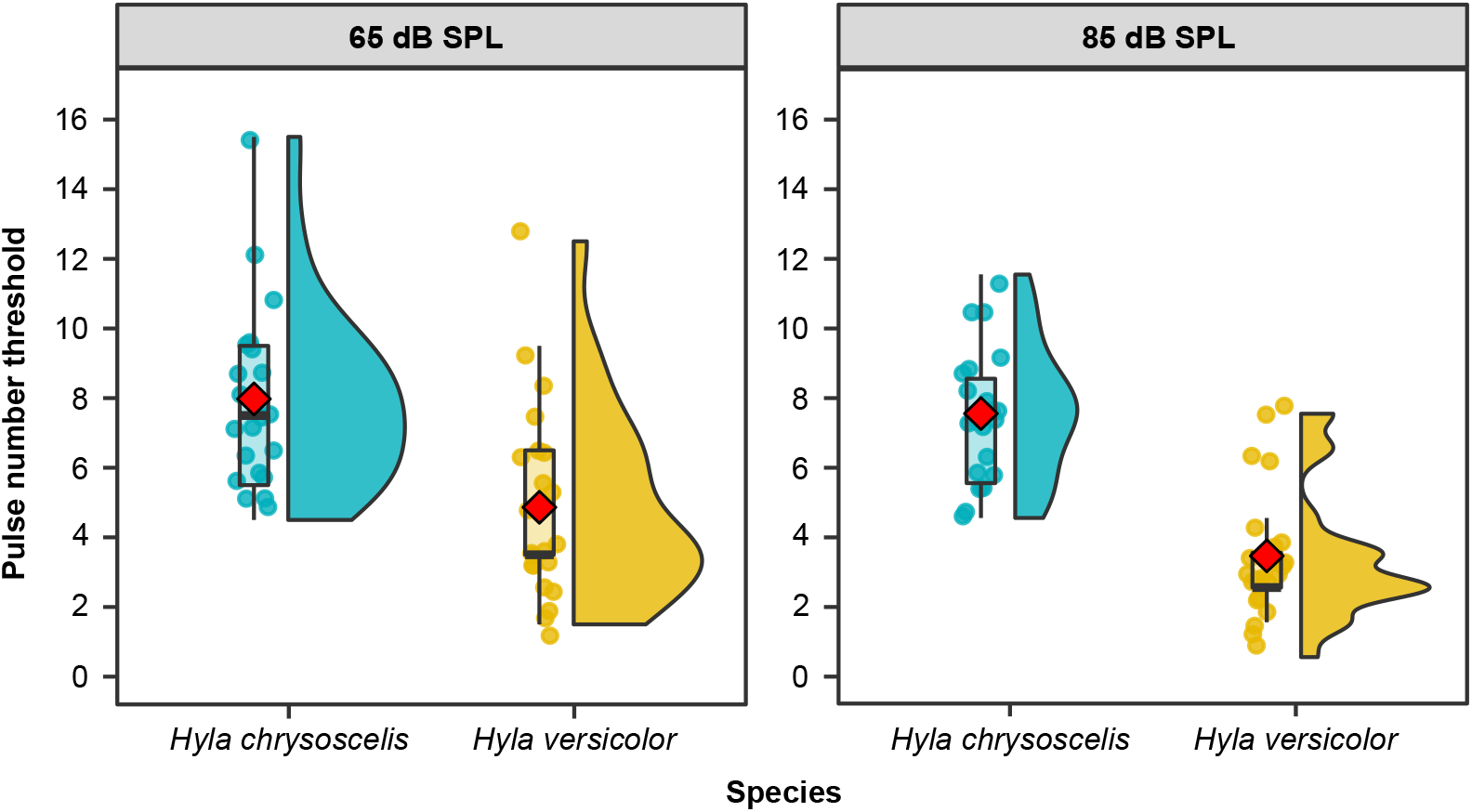
Behavioral pulse number thresholds. Raincloud plots that combine box plots, raw jittered data, and split-half violins depict behavioral pulse number thresholds obtained from *H. chrysoscelis* (blue) and *H. versicolor* (yellow). In the box plots, the lower and upper boundaries of the box indicate 25^th^ and 75^th^ percentile, respectively. The whiskers below and above the box indicate the smallest value within 1.5 times interquartile range below 25^th^ percentile and the largest value within 1.5 times interquartile range above 75^th^ percentile, respectively. The black line inside the box indicates the median and the red point indicates the mean. The transparent points superimposing the box plot depict individual data points and the split-half violin plots depict the kernel density of data. The left and right panels depict pulse number thresholds determined at 65 dB and 85 dB SPL, respectively. On average, behavioral pulse number threshold of *H. chrysoscelis* was higher than that of *H. versicolor*. Additionally, threshold obtained at 65 dB SPL was also, on average, higher than that obtained at 85 dB SPL.

**Table 1.** Output of a linear mixed-effect model fitted to compare behavioral pulse number thresholds between the two species and between the two signal amplitudes.

|  |  | Estimate | Standard Error | t value | P |
| --- | --- | --- | --- | --- | --- |
| (Intercept) |  | 0.49343 | 0.04018 | 12.280 | << 0.001 |
| Species | *Hyla chrysoscelis* vs *Hyla versicolor* | 0.33125 | 0.05125 | 6.464 | << 0.001 |
| Signal Amplitude | 65 dB vs 85 dB SPL | 0.09245 | 0.03583 | 2.580 | 0.0133 |

### Neural Pulse Number Thresholds

Neural pulse number thresholds were higher for *H. chrysoscelis* than for *H. versicolor* in both ICN populations (Fig. 3). Regression analysis showed that neural pulse number thresholds of *H. chrysoscelis* were significantly higher (by a factor of 1.70; *β* = 0.23, *p* < 0.001; Table 2) than those of *H. versicolor* after controlling for differences between band-pass and band-suppression ICNs. Also, pulse number thresholds of band-pass ICNs were significantly higher (by a factor of 2.63; *β* = 0.42, *p* < 0.001; Table 2) than those of band-suppression ICNs after controlling for species differences. The mean pulse number threshold of band-pass ICNs in *H. chrysoscelis* was 6.62 ± 1.27 (Median = 6.0, IQR = 5.0-7.25, N = 16) and that in *H. versicolor* was 3.94 ± 0.64 (Median = 4.0, IQR = 3.0-5.0, N = 17). The mean pulse number threshold of band-suppression ICNs in *H. chrysoscelis* was 2.75 ± 0.84 (Median = 2.0, IQR = 2.0-3.25, N = 12) and that in *H. versicolor* was 1.56 ± 0.66 (Median = 1.0, IQR = 1.0-2.0, N = 9).

**Fig. 3.**
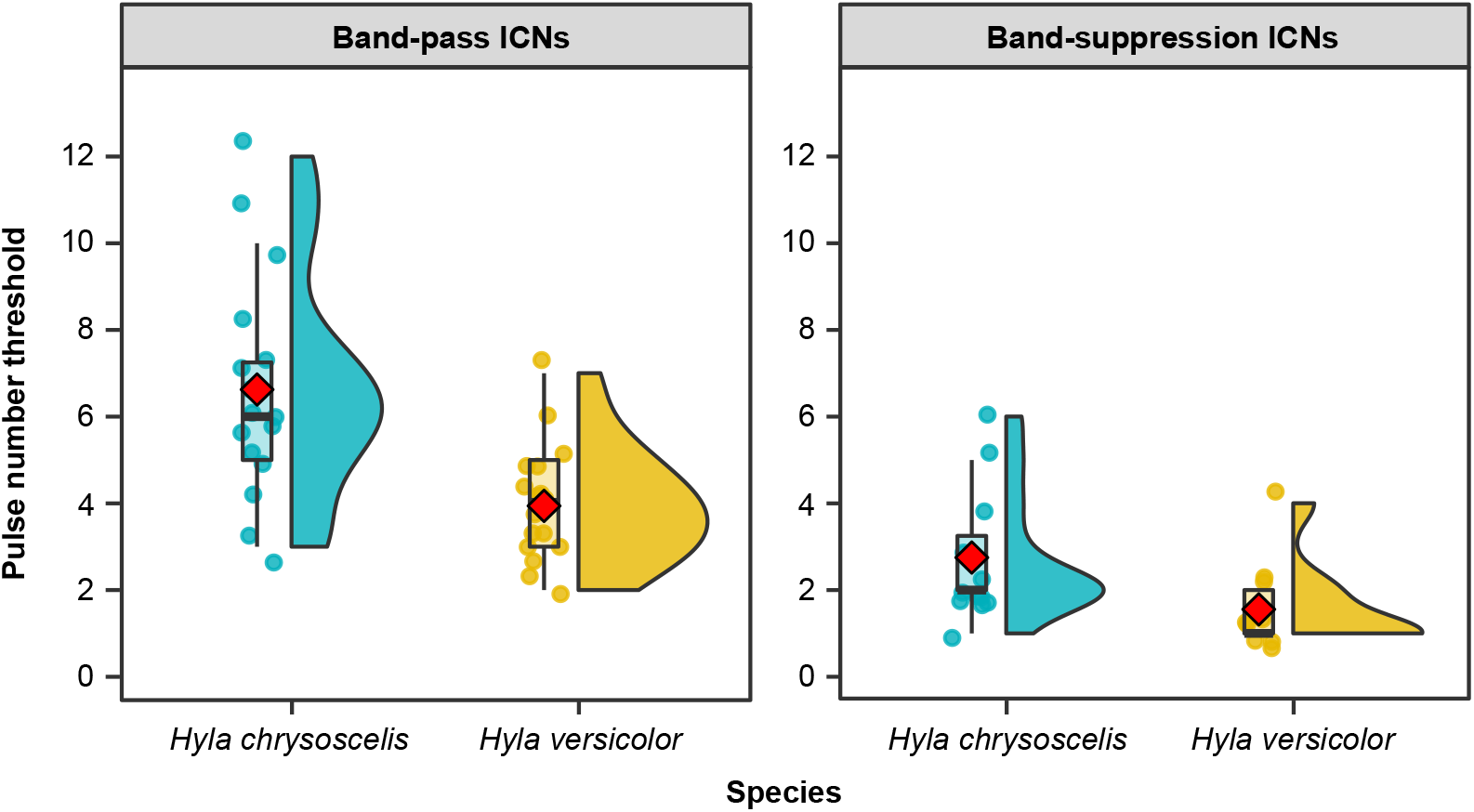
Pulse number thresholds of ICNs. Raincloud plots depicting pulse number thresholds of ICNs in *Hyla chrysoscelis* (blue) and *Hyla versicolor* (yellow). Refer to Fig. 2 for explanation of raincloud plots and its individual components. The left and the right panels depict pulse number thresholds determined from band-pass and band-suppression ICNs, respectively. On average, neural pulse number threshold of *H. chrysoscelis* was higher than that of *H. versicolor*. Also, the average pulse number threshold of band-pass ICNs was higher than band-suppression ICNs.

**Table 2.** Output of a linear regression model fitted to compare neural pulse number thresholds between the two species and between the two ICN populations.

|  |  | Estimate | Standard Error | t value | P |
| --- | --- | --- | --- | --- | --- |
| (Intercept) |  | 0.14694 | 0.04923 | 2.985 | < 0.01 |
| Species | *Hyla chrysoscelis* vs *Hyla versicolor* | 0.23182 | 0.05014 | 4.624 | << 0.001 |
| ICN populations | Band-pass vs Band-suppression ICNs | 0.41775 | 0.05139 | 8.129 | << 0.001 |

### Comparison Between Behavioral and Neural Pulse Number Thresholds

In both species, behavioral pulse number thresholds were similar to the pulse number thresholds of band-pass ICNs, but higher than those of band-suppression ICNs (Fig. 4). Within *H. chrysoscelis*, behavioral pulse number thresholds were not significantly different from the pulse number thresholds of band-pass ICNs (*β* = 0.09, *p* = 0.106), but they were significantly higher (by a factor of 3.09; *β* = 0.49, *p* < 0.001) than the thresholds of band-suppression ICNs (Table 3). Similarly, behavioral pulse number thresholds in *H. versicolor* were also not significantly different from the pulse number thresholds of band-pass ICNs (*β* = 0.005, *p* = 0.941), but they were significantly higher (by a factor of 2.69; *β* = 0.43, *p* < 0.001) than the thresholds of band-suppression ICNs (Table 3).

**Fig. 4.**
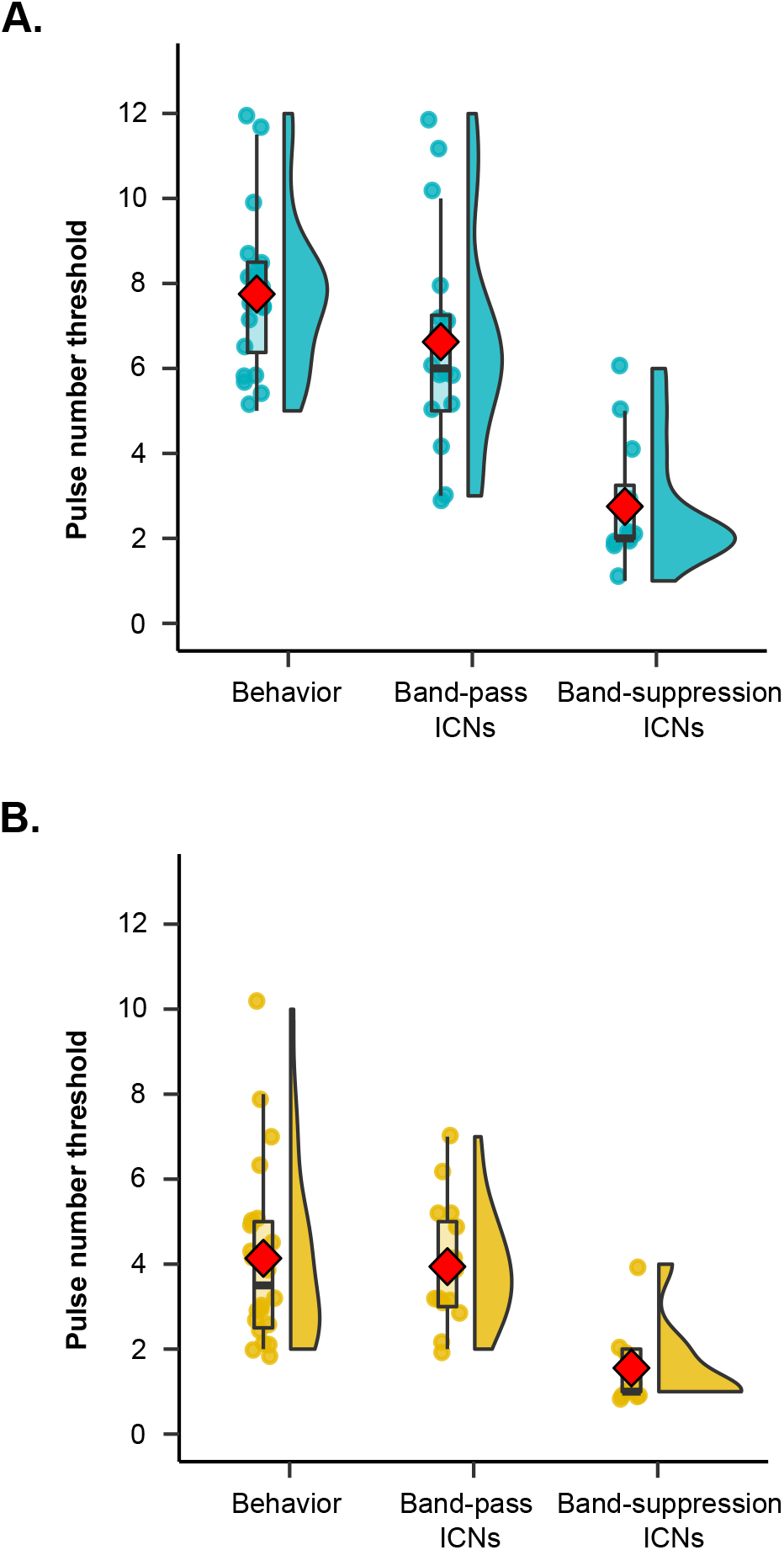
Comparison between behavioral and neural pulse number thresholds. Raincloud plots depicting pulse number thresholds obtained from behavioral measurements, band-pass ICNs and band-suppression ICNs in (A) *Hyla chrysoscelis* (blue) and (B) *Hyla versicolor* (yellow) subjects. Refer to Fig. 2 for explanation of raincloud plots and its individual components. In both species, behavioral thresholds closely match the thresholds obtained from band-pass ICNs, but not band-suppression ICNs.

**Table 3.** Output of two linear regression models fitted separately for each species to compare behavioral and neural pulse number thresholds obtained from each ICN population. The reference level in each model is behavioral pulse number threshold.

|  | Estimate | Standard Error | t value | P |
| --- | --- | --- | --- | --- |
| ***Hyla chrysoscelis*** |  |  |  |  |
| (Intercept) | 0.87798 | 0.03593 | 24.438 | << 0.001 |
| Band-pass ICNs | -0.08887 | 0.05389 | -1.649 | 0.106 |
| Band-suppression ICNs | -0.48936 | 0.05967 | -8.341 | << 0.011 |
| ***Hyla versicolor*** |  |  |  |  |
| (Intercept) | 0.56705 | 0.04086 | 13.876 | << 0.001 |
| Band-pass ICNs | 0.00461 | 0.06190 | 0.074 | 0.941 |
| Band-suppression ICNs | -0.43325 | 0.07584 | -5.713 | << 0.001 |

## DISCUSSION

### Interval-Counting Neurons Decode Temporal Information for Species Recognition

Many closely related species use the temporal features of signals to correctly identify mates of their own species, but less is known about the neural processes that generate differences in temporal pattern recognition between species. We tested the hypothesis that neurons in the frog midbrain that count inter-pulse intervals specified by conspecific pulse rates mediate acoustic species recognition in a cryptic species complex. Within each species of gray treefrog, we found a close correspondence between the threshold number of pulses required to elicit both positive phonotaxis from females and action potentials from ICNs exhibiting band-pass tuning for conspecific pulse rates. More importantly, both behavioral and neural pulse number thresholds differed in parallel between the two cryptic species: the average pulse number thresholds obtained behaviorally and from band-pass ICNs ranged between 6 and 8 pulses in *H. chrysoscelis* and between 3 and 5 pulses in *H. versicolor*. In contrast, the average pulse number thresholds of band-suppression ICNs were similar between the two species and relatively lower (ranging between 1 and 3 pulses) compared with thresholds determined behaviorally and from band-pass ICNs. The agreement between the pulse number thresholds of females and band-pass ICNs observed *within* each species, combined with the consistent differences in behavioral and neural thresholds observed *between* species, strongly suggests band-pass ICNs play an important functional role in behaviors that depend on correctly decoding temporal information relevant to species identity.

The present study, which integrated behavioral and neurophysiological investigations of a cryptic species complex, lends considerable support to the emerging view that interval-counting neurons play key roles in decoding information about species identity in frogs. In the Pacific treefrogs, *Pseudacris regilla*, for example, Rose and Brenowitz (1997) showed in a field playback experiment that advertisement calls with pulse rates between 80 to 120 pulses/s evoked aggressive behavior from males. Pulse sequences in which intervals alternated between those characteristic of advertisement or encounter calls were treated as being the latter; at least 4 consecutive pulses with intervals characteristic of those in advertisement calls were required to significantly elevate aggressive thresholds in response to playbacks of advertisement calls. A neurophysiological study later demonstrated that many ICNs in this species are, in fact, selective for the species-specific pulse rates typical of their advertisement calls (Edwards and Rose, 2003; Edwards et al., 2007). Similarly, behavioral and neurophysiological studies of gray treefrogs have shown that females of *H. chrysoscelis* and *H. versicolor* prefer advertisement calls with conspecific pulse rates of approximately 30-60 pulses/s and 10-30 pulses/s, respectively, and that a large proportion of IC neurons, including ICNs, found in these two species are also tuned to conspecific pulse rates (Bush et al., 2002; Diekamp and Gerhardt, 1995; Rose et al., 1985; Rose et al., 2015; Schul and Bush, 2002). Schwartz et al. (2010) demonstrated that females of *H. versicolor* discriminated against previously attractive calls when the species-typical inter-pulse interval was artificially increased or decreased with a call. This outcome was interpreted as a behavioral correlate of the resetting of interval counting by ICNs in response to anomalous inter-pulse intervals (Edwards et al., 2002). Our estimates of behavioral pulse number thresholds in gray treefrog corroborate related data from previous studies suggesting that female of *H. versicolor* potentially respond behaviorally to calls having fewer pulses (e.g., 3 to 6; Bush et al., 2002) compared with females of *H. chrysoscelis* (e.g., 6 to 9; Vélez and Bee, 2011). Our study extends these previous investigations by showing that a species difference in the minimum number of consecutive pulses required to evoke a behavioral response to conspecific calls corresponds closely with a parallel species difference in the neural responses of a specific population of midbrain neurons, namely band-pass tuned ICNs, that function in decoding species-specific temporal information.

### Proximate Explanations for the Species Differences in Pulse Number Thresholds

At a proximate level, it is important to consider two potential explanations for the observed species differences in pulse number thresholds. On the one hand, a species difference in pulse number thresholds would be expected any time two species have different pulse rates but integrate information over similar time windows: species with faster pulse rates would necessarily have higher pulse number thresholds, consistent with our observations. However, the two gray treefrogs do not appear to share a common integration time. When the observed pulse number thresholds are instead expressed as threshold integration times — i.e., the minimum call duration required to elicit a response — *H. chrysoscelis* had threshold integration times that were lower (by a factor of 0.71 for behavioral responses and 0.76 for neural responses) compared with *H. versicolor* (see Supplementary Materials). Hence, the data do not strongly support an explanation of species differences in pulse number thresholds based on common integration times but differing pulse rates. Instead, we believe the observed species difference more likely reflects underlying differences in the cellular and circuit mechanisms responsible for “counting” individual pulses.

Experimental data from whole-cell recordings of ICNs show that the temporal dynamics of fast inhibition and slower build-up of rate-dependent excitation shape the pulse number thresholds of ICNs (Edwards et al., 2007; Rose, 2014). Hence, a change in the balance between the time course of excitation and inhibition events potentially leads to differences in pulse number thresholds (Edwards et al., 2007). A computational model built to explain the experimental data provides further insights into the potential mechanisms of interval counting (Naud et al., 2015), and can be used to generate testable hypotheses about factors that might contribute to the species differences in pulse number thresholds observed in the present study. According to this model, ICNs receive excitatory input via AMPA- and NMDA-type receptors and inhibitory input from long-interval neurons (LINs). When a pulsatile stimulus having an optimal pulse rate is presented, the initial few pulses activate LINs, which provide strong inhibition to ICNs. This inhibition initially counteracts excitation, but with additional pulses, LINs themselves receive inhibitory input via a feed-forward mechanism. The release of inhibition allows the excitatory inputs to cause depolarization of ICNs, which is further augmented by the recruitment of NMDA-type voltage-dependent conductances. This “dis-inhibitory” network model predicts that the pulse number threshold of ICNs primarily depends on the number of excitatory inputs that synapse onto ICNs, the decay time of those excitatory inputs, and the time course of rate-dependent depression of inhibitory inputs (Naud et al., 2015). For example, increasing the strength and slowing the rate of depression of inhibition would be expected to increase pulse number thresholds; reducing the strength excitatory inputs and decreasing the time course of excitation could lead to a similar increase. We hypothesize that these differences can explain the species differences reported here. Whether such differences might arise directly as a result of polyploid speciation, from selection for greater species isolation following polyploidization, or both, could be investigated by generating artificial polyploids (Keller and Gerhardt, 2001; Tucker and Gerhardt, 2012). Our preliminary recordings from autotriploids of *H. chrysoscelis* support the hypothesis that polyploidy alone can shift the temporal selectivity of midbrain neurons towards slower pulse rates and longer rise times, as seen in *H. versicolor* (unpublished data).

### Functional Role of Band-Suppression ICNs

Unlike band-pass ICNs, which selectively respond to relatively narrow ranges of pulse rates, band-suppression ICNs show a selectively reduced response to intermediate pulse rates, relative to spike rates for slower or faster pulse rates. In Pacific treefrogs, the pulse rates of conspecific advertisement calls are able to drive responses in band-suppression ICNs (Edwards and Rose, 2003; Rose et al., 2015). Hence, they might play an important role in advertisement call recognition. However, our finding that the pulse number thresholds of band-suppression ICNs are dissimilar to behavioral pulse number threshold casts doubt on this notion. An alternate hypothesis is that band-suppression ICNs play some role in the recognition of aggressive calls. This conjecture is based on reports that band-suppression ICNs in Pacific treefrogs are tuned to both fast pulse rates (~90 pulses/s) typical of advertisement calls and to slower pulse rates (~25 pulses/s) more typical of aggressive calls (Edwards and Rose, 2003, Rose and Brenowitz, 2002). In gray treefrogs, aggressive calls typically lack the stereotypical pulsatile structure found in advertisement calls (Reichert and Gerhardt, 2014). However, aggressive calls consist of long-duration pulses, which are effective stimuli for band-suppression cells. The potential role of band-suppression ICNs in aggressive call recognition could be tested in future studies by recording the activity of band-suppression ICNs in response to conspecific aggressive calls.

### Relation to Previous Work on Neural Mechanisms Underlying Species-Specific Behavior

In this study, similarities in behavioral and neural pulse number thresholds within species, and parallel differences between species, suggest temporal processing by a subset of midbrain neurons plays an important role in guiding species-specific call recognition. This finding adds to a growing body of experimental and modeling studies investigating the neural mechanisms underlying temporal pattern recognition in closely related species (Hennig, 2003; Hennig et al., 2014; Triblehorn and Schul, 2009). In two congeneric field crickets, *Teleogryllus oceanicus* and *T. commodus*, for example, females differ in their selectivity for the temporal features of male calling song that promote species recognition. In the former, species recognition is based on selectivity for the rate of pulses, whereas in the latter, it is based on selectivity for the duration of pulses (Hennig, 2003). Using a cross-correlation signal analysis method, Hennig (2003) suggested that a simple change in the timescale of oscillatory properties of neurons or neuronal networks during speciation can explain the observed differences in temporal pattern selectivity between the two sister species. Taken together, the present study and previous reports suggest that differences in the response properties of key elements of neural circuits shape divergence in sound pattern recognition in ways that facilitate premating reproductive isolation among closely related species. These findings complement research demonstrating that subtle differences in vocal circuits among closely related species of frogs underlie the production of temporally distinct sound patterns (Barkan et al., 2017; Barkan et al., 2018). Together, this work on acoustically signaling insects and frogs supports the general notion that evolutionarily functional differences in behavior between species can result from small changes in homologous neural pathways (Katz, 2011; Katz and Harris-Warrick, 1999).

### Ethics

The behavioral procedures were approved by the University of Minnesota’s Institutional Animal Care and Use Committee (2001-37746A) and adhered to the ‘Guidelines for the treatment of animals in behavioural research and teaching’, jointly published by the Animal Behavior Society and the Association for the Study of Animal Behaviour. The neurophysiological procedures were approved by The University of Utah’s Institutional Animal Care and Use Committee (19-11-008) and were conducted in compliance with the National Institutes of Health guidelines.

## Acknowledgements

We thank all members of the Bee Lab 2019, especially the undergrads, for assistance in collecting and testing animals, the Minnesota Department of Natural History for permission to collect gray treefrogs, and the Ramsey County Department of Parks and Recreation for after-hour access to the field site. We are also grateful to Kyphoung Luong of the Rose Lab for his help in collecting neurophysiological data.

## Author Contribution

S.G and M.A.B designed the behavioral experiments. S.G. conducted the behavioral experiments and analyzed the data. R.K.A and G.J.R designed and conducted the neurophysiological experiments. S.G and G.J.R analyzed the neural results. S.G and M.A.B wrote the manuscript. All authors discussed the results and edited the manuscript.

## Data Availability

Data is publicly available at https://doi.org/10.13020/6c7h-4b35

