## Supplementary Materials for "Neural Basis of Acoustic Species Recognition in a Cryptic Species Complex"

**Recording and Analysis of *Hyla versicolor* Advertisement Calls**

We recorded and analyzed the advertisement calls of *H. versicolor* using methods described elsewhere to record and analyze the calls of *H. chrysoscelis* (Ward et al., 2013). Briefly, during the gray treefrog breeding season (from May to June) of 2006, the advertisement calls of 14 *H. versicolor* males were recorded at night between 2200 and 0100 hours in wetlands and ponds located in east-central Minnesota, U.S.A. (Tamarack Nature Center, Ramsey County, 45°06’12’’N, 93°02’27’’W; Lake Maria State Park, Wright County, 45°19’13’’N, 93°56’37’’W). A Sennheiser ME66 microphone and K6 power supply (Sennheiser USA, Old Lyme, CT, U.S.A.) connected to a Marantz PMD670 recorder (Marantz Professional, Cumberland, RI, USA) (44.1 kHz sampling rate, 16-bit resolution) were used to record the calls. The tip of the microphone was directed towards the focal male and positioned approximately 1 m away. A quick-reading Miller & Weber thermometer (Avinet Inc., Dryden, NY, USA) was used to measure the water and wet-bulb air temperatures (to the nearest 0.1°C) at the calling location of each male.

We used the automatic call and pulse detection and measurement functions of SoundRuler (Bee, 2004; Gridi-Papp, 2007) to segment recordings into individual calls composed of individual pulses (which we manually verified) and to determine the values of 11 temporal properties (time resolution = 5.8 ms) and 3 spectral properties (FFT length = 256, 90% overlap, 172.3 Hz frequency resolution). The following temporal properties were determined for each call. Call duration was measured as the time (in ms) between the onset of the first pulse of the offset of the last pulse in a call. Because pulse number carries essential information in gray treefrogs (Bush et al., 2002; Ward et al., 2013; Welch et al., 1998), call duration was also measured as the total number of pulses/call. Call period (in s) was measured as the time between the onset of the first pulse in a call to the onset of the first pulse in the next call. Pulse period was determined as the mean duration (in ms) from the onset of one pulse to onset of the next pulse in a call, and it was computed by averaging over all pulse periods in each call. The pulse rate (pulses/s) of each call was computed by taking the inverse of the mean pulse period and multiplying it by 1000 ms. Pulse duration was determined as the mean duration (in ms) from the onset of one pulse to the offset of the same pulse, and it was computed by averaging over all pulse durations in each call. Pulse duty cycle was computed as a proportion by dividing the mean pulse duration by the mean pulse period for each call. Mean values of pulse rise time and pulse fall time (in ms), which are the durations from the onset of one pulse to the point where the amplitude reaches its maximum and the duration from the point of maximum amplitude to the offset of a call, respectively, were also computed for each call. Similarly, 50% pulse rise time and 50% pulse fall time (in ms) were calculated, respectively, by measuring the time between the onset of one pulse and the point where it reached 50% of its maximum amplitude and the time between the maximum amplitude and the subsequent point where the amplitude had decreased to 50% of its maximum value. For each pulse, we also measured the frequencies (in kHz) of the first and second harmonics, which correspond to the fundamental frequency and dominant frequency, respectively, and the amplitude of the first harmonic relative to that of the second harmonic (in dB; hereafter “relative amplitude”). These spectral properties were then averaged to determine the mean values for each individual call. We then computed a mean value of all 14 properties for each individual male before computing the mean (± SD), median (interquartile range), and range for each call property over the entire sample of 14 individuals (Table S1).

The well-known influence of ambient temperature on the properties of frog calls was taken into account by adjusting all values to a common temperature of 20°C following Platz and Forester (1988). A temperature of 20°C is commonly used to standardize measures of gray treefrog calls and for conducting behavioral experiments. For each male, the water temperature or wet-bulb air temperature was selected based on the frog’s calling site. For example, if the frog was sitting on the surface of the water while calling, the water temperature was used. But if it was sitting on emergent vegetation, then the wet-bulb air temperature was used for the analysis. The mean (± SD), median (interquartile range), and range of each temperature-corrected call parameter computed over the entire sample of 14 individuals are reported in Table S2.

**Threshold Integration Times**

To assess whether the observed species differences in pulse number thresholds could be explained by the two species having different pulse rates but integrating information over similar time windows, we converted the observed pulse number thresholds into threshold integration times, which we defined as the minimum stimulus duration required to elicit a behavioral or neural response. Conversions were based on the known pulse durations and pulse periods in the synthetic stimuli used for each species.

Following the methods used for pulse number threshold analyses, behavioral and neural threshold integration times were analyzed using a linear mixed effect model and a linear regression model, respectively. Threshold integration times were $\log_{10}$ transformed to meet the assumptions of normality and homogeneity of variance and all estimates were subsequently back-transformed to report the effect sizes. A significance criterion of $\alpha = 0.05$ was used for hypothesis testing.

Threshold integration times were lower in *H. chrysoscelis* than in *H. versicolor* (Supp. Fig. 1). For behavioral measurements, the threshold integration time of *H. chrysoscelis* was significantly lower (by a factor of 0.71; $\beta = -0.15$, $p = 0.007$) than that of *H. versicolor* after controlling for the effects of signal amplitude. The threshold integration time determined at 65 dB SPL was significantly higher (by a factor of 1.23; $\beta = 0.09$, $p = 0.013$) than that determined at 85 dB SPL after controlling for species differences. For neural measurements, the threshold integration time of band-pass ICNs in *H. chrysoscelis* was 0.76 times lower than that of band-pass ICNs in *H. versicolor*, although, this difference was not quite statistically significant ($\beta = -0.12$, $p = 0.07$).

**Table S1.** Descriptive statistics summarizing acoustical properties of *Hyla versicolor* advertisement calls from Minnesota prior to temperature correction (*N* =14). The temperature range at which the calls were recorded was 10.2 – 21.8°C.

| Call Parameter | Mean ± SD | Median (IQR) | Range |
|---|---|---|---|
| Call duration (ms) | 838.8 ± 282.9 | 802.3 (583.2 – 1103.3) | 355.6 – 1261.1 |
| Call duration (number of pulses) | 16.3 ± 2.3 | 16.1 (14.1 – 18.9) | 13.0 – 19.7 |
| Call period (s) | 5.9 ± 1.9 | 5.5 (4.2 – 7.3) | 3.2 – 9.4 |
| Fundamental frequency (kHz) | 1.2 ± 0.1 | 1.2 (1.1 – 1.2) | 1.1 – 1.5 |
| Dominant frequency (kHz) | 2.5 ± 0.2 | 2.4 (2.3 – 2.5) | 2.2 – 3.0 |
| Relative amplitude (dB) | -15.0 ± 2.3 | -14.7 (-16.3 – -13.3) | -19.2 – -11.6 |
| Pulse period (ms) | 61.1 ± 19.4 | 49.1 (46.0 – 82.6) | 41.2 – 94.1 |
| Pulse rate (pulses/s) | 17.9 ± 5.0 | 20.6 (12.1 – 21.8) | 10.7 – 24.3 |
| Pulse duration (ms) | 22.4 ± 5.2 | 20.2 (18.3 – 28.5) | 16.2 – 30.9 |
| Pulse duty cycle | 0.3 ± 0.06 | 0.3 (0.27 – 0.33) | 0.16 – 0.36 |
| Pulse rise time (ms) | 14.5 ± 4.1 | 13.0 (11.2 – 17.6) | 10.9 – 22.0 |
| 50% pulse rise (ms) | 8.3 ± 2.2 | 7.7 (6.8 – 10.2) | 6.0 – 13.2 |
| Pulse fall time (ms) | 7.4 ± 1.8 | 7.2 (6.0 – 8.7) | 4.5 – 10.9 |
| 50% pulse fall (ms) | 4.2 ± 1.4 | 4.1 (3.0 – 5.3) | 2.4 – 7.5 |

**Table S2.** Descriptive statistics summarizing the temperature-corrected (to 20°C) acoustical properties of *Hyla versicolor* advertisement calls from Minnesota (*N* =14). The temperature range at which the calls were recorded was 10.2 – 21.8°C.

| Call Parameter | Mean ± SD | Median (IQR) | Range |
|---|---|---|---|
| Call duration (ms) | 643.2 ± 134.0 | 652.2 (561.9 – 697.4) | 383.9 – 867.3 |
| Call duration (number of pulses) | 16.0 ± 2.8 | 15.6 (13.9 – 18.4) | 12.3 – 19.3 |
| Call period (s) | 4.5 ± 1.0 | 4.5 (4.0 – 5.0) | 2.7 – 6.7 |
| Fundamental frequency (kHz) | 1.2 ± 0.09 | 1.2 (1.17 – 1.25) | 1.1 – 1.5 |
| Dominant frequency (kHz) | 2.5 ± 1.80 | 2.4 (2.3 – 2.5) | 2.2 – 3.0 |
| Relative amplitude (dB) | -13.9 ± 1.9 | -14.0 (-14.9 – -12.8) | -17.9 – -10.1 |
| Pulse period (ms) | 49.5 ± 13.0 | 48.5 (39.6 – 58.9) | 29.4 – 72.7 |
| Pulse rate (pulses/s) | 21.1 ± 2.9 | 20.8 (19.0 – 23.6) | 16.6 – 25.7 |
| Pulse duration (ms) | 19.6 ± 3.9 | 19.8 (16.2 – 23.3) | 13.1 – 25.2 |
| Pulse duty cycle | 0.3 ± 0.05 | 0.3 (0.28 – 0.35) | 0.18 – 0.38 |
| Pulse rise time (ms) | 12.6 ± 3.3 | 11.9 (10.2 – 14.1) | 6.9 – 18.4 |
| 50% pulse rise (ms) | 7.5 ± 1.9 | 7.2 (6.0 – 8.7) | 4.7 – 11.6 |
| Pulse fall time (ms) | 6.6 ± 1.5 | 6.5 (5.7 – 7.1) | 3.5 – 9.2 |
| 50% pulse fall (ms) | 3.3 ± 1.0 | 3.4 (2.5 – 4.3) | 1.4 – 4.7 |

**Table S3**. ANOVA output comparing two linear mixed effects models fitted for analyzing behavioral pulse number thresholds.

| **Model** | **AIC** | **BIC** | **Log likelihood** | **Deviance** | $\chi^2$ | **Df** | ***P*** |
|---|---|---|---|---|---|---|---|
| Thresholds ~ Species + Amplitude + (1\|Subject ID) | -23.77 | -11.50 | 16.89 | -33.77 | | | |
| Thresholds ~ Species + Amplitude + Species * Amplitude + (1\|Subject ID) | -25.20 | -10.47 | 18.60 | -37.20 | 3.42 | 1 | 0.06 |

**Table S4**. ANOVA output comparing two linear regressions fitted for analyzing neural pulse number thresholds.

| **Model** | **Residue Df** | **Model Df** | **Sum of squares** | **Df** | **F value** | ***P*** |
|---|---|---|---|---|---|---|
| Thresholds ~ Species + ICN population | 51 | 1.7161 | | | | |
| Thresholds ~ Species + ICN population + Species * ICN population | 50 | 1.7116 | 0.0044241 | 1 | 0.1292 | 0.7207 |

**Table S5**. Output from a set of linear regression models comparing the behavioral pulse number thresholds determined at 65 and 85 dB SPL, respectively with neural pulse thresholds of the two populations of ICNs in *Hyla chrysoscelis*.

| | **Estimate** | **Standard Error** | **t value** | ***P*** |
|---|---|---|---|---|
| **Reference level: Behavior determined at 65 dB SPL** | | | | |
| (Intercept) | 0.88147 | 0.03703 | 23.806 | << 0.01 |
| Band-pass ICNs | -0.09236 | 0.05631 | -1.640 | 0.108 |
| Band-suppression ICNs | -0.49285 | 0.06140 | -8.026 | << 0.01 |
| **Reference level: Behavior determined at 85 dB SPL** | | | | |
| (Intercept) | 0.86073 | 0.03575 | 24.079 | << 0.001 |
| Band-pass ICNs | -0.07161 | 0.05436 | -1.317 | 0.194 |
| Band-suppression ICNs | -0.47210 | 0.05928 | -7.964 | << 0.001 |

**Table S6**. Output from a set of linear regression models comparing the behavioral pulse number thresholds determined at 65 and 85 dB SPL, respectively with neural pulse thresholds of the two populations of ICNs in *Hyla versicolor*.

| | **Estimate** | **Standard Error** | **t value** | ***P*** |
|---|---|---|---|---|
| **Reference level: Behavior determined at 65 dB SPL** | | | | |
| (Intercept) | 0.61747 | 0.04641 | 13.305 | << 0.001 |
| Band-pass ICNs | -0.04582 | 0.07029 | -0.652 | 0.518 |
| Band-suppression ICNs | -0.48368 | 0.08613 | -5.615 | << 0.001 |
| **Reference level: Behavior determined at 85 dB SPL** | | | | |
| (Intercept) | 0.46183 | 0.04781 | 9.661 | << 0.001 |
| Band-pass ICNs | 0.10982 | 0.07241 | 1.517 | 0.13632 |
| Band-suppression ICNs | -0.32804 | 0.08872 | -3.697 | < 0.001 |

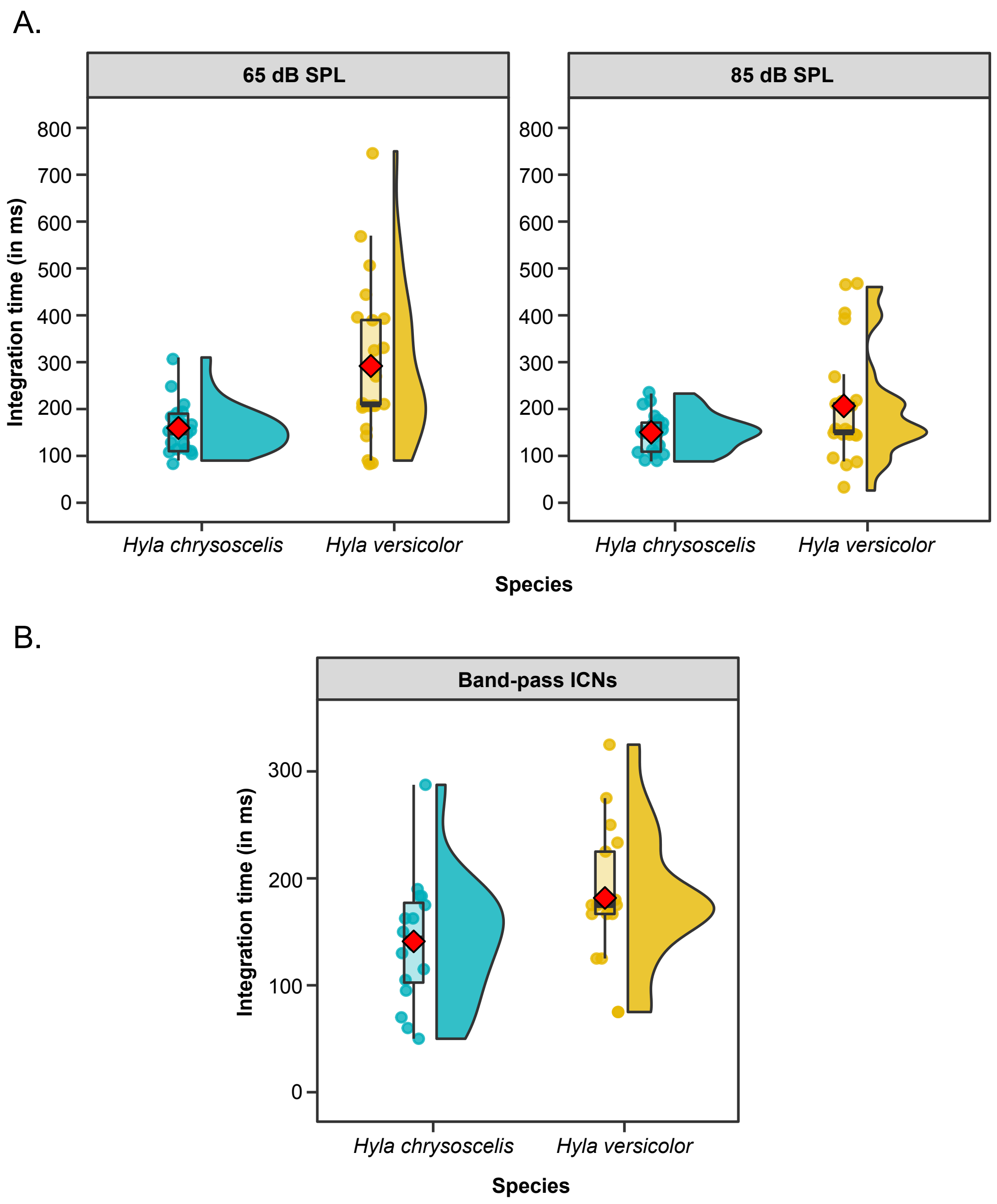


**Fig. S1. Threshold integration times.** Raincloud plots depicting threshold integration time. (A) Behavioral threshold integration times for female *H. chrysoscelis* (blue) and *H. versicolor* (yellow) determined at 65 dB SPL (left) and 85 dB SPL (right). (B) Neural threshold integration times of band-pass ICNs in *H. chrysoscelis* and *H. versicolor*.
